# Nanoscale photobiotinylation, pulldown and sequencing of region-specific DNA from intact cells

**DOI:** 10.1101/2023.12.04.569951

**Authors:** Thomas C. Roberts, Max Kushner, Jack C. Crowley, Abdullah Ozer, Juan Wang, Judhajeet Ray, John T. Lis, Warren R. Zipfel

## Abstract

Femto-seq is a novel nanoscale optical method that can be used to obtain DNA sequence information from targeted regions around a specific locus or other nuclear regions of interest. Two-photon excitation is used to photobiotinylate femtoliter volumes of chromatin within the nucleus, allowing for subsequent isolation and sequencing of DNA, and bioinformatic mapping of any nuclear region of interest in a select set of cells from a heterogenous population.

---

Spatio-temporal changes in nuclear chromatin organization are fundamental to understanding how the nucleus functions. The detailed structural organization of the genome has become clearer over the past few decades with the development of high-resolution microscopy coupled with multiplexed FISH and in situ genome sequencing, sequencing-based methods such as Hi-C and its derivatives, and new computational approaches^1^.

Here we present a new approach designed to look at the genomic sequences located within a region of interest from user-selected cells of interest. The method, called Femto-seq (Figure 1), combines 3D localized two-photon excitation dependent photo-biotinylation in targeted nuclear volumes as small as a cubic micron (femtoliter) followed by chromatin isolation and DNA sequencing. The nuclear volumes from which sequence information can be obtained can be as small as the two-photon point spread function (PSF) of the microscope (Fig. 1D-F). Upon light membrane permeabilization, our optimized photoactivatable DNA crosslinker with a biotin affinity tag^2^ (AP3B – Fig 1A) enters cells and intercalates into DNA. Using confocal microscopy, the fluorescently labeled regions of interest (ROIs) are located and two-photon excitation at 700 nm is used to photoactivate the crosslinker within these regions (Fig. 1G). ROIs in the nucleus are assigned based on 3D confocal imaging of fluorescence markers. ROIs can be labeled gene loci, nuclear bodies or other distinguishable structures, and since identification is imaging based, the user can select which cells are targeted. For example, a sub-set may be targeted based on cell state or expression of a specific marker. After crosslinking, cells are washed to remove non-crosslinked AP3B. Chromatin is extracted from the pre-washed cells, and then sheared and purified using the biotin tag. After crosslink reversal and adapter ligation, the captured chromatin fragments are sequenced. Sequencing data can provide information about the spatial association between regulatory sequences and the targeted gene loci, or sequences associated with certain sub-nuclear compartments.

**Figure 1.**
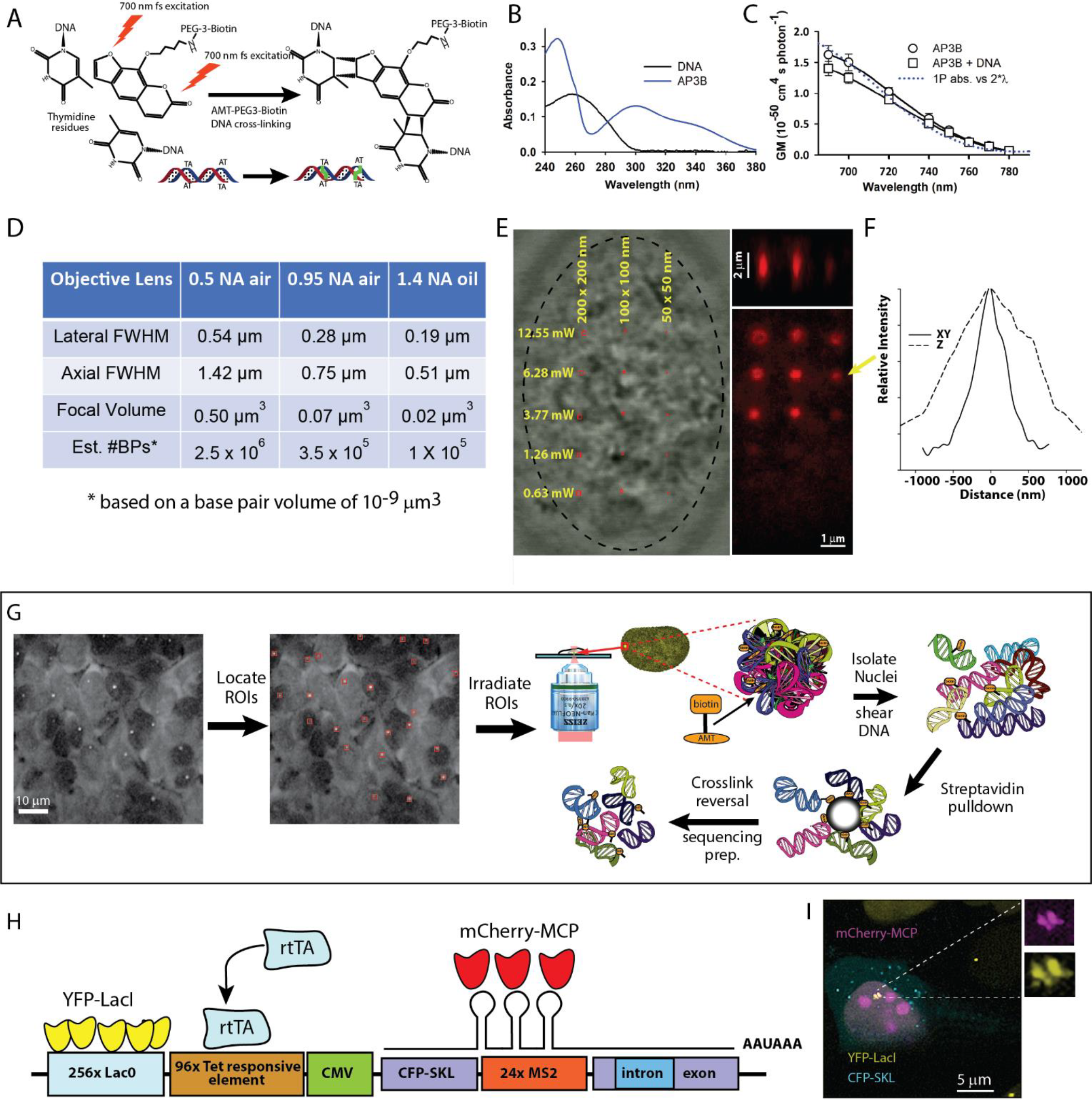
Targeted nanoscale photobiotinylation using Femto-seq. **A**. 4’-aminomethyltrioxsalen (4’-AMT) is a psoralen family DNA cross-linker, which forms covalent bonds primarily with thymidine residues when excited by UV or UV-like two-photon (2P) irradiation. The photoactivatable reagent used is 4’-Aminomethyltrioxsalen-PEG3-Biotin (AP3B), which enables covalent attachment of biotin to DNA. **B**. Absorbance spectra of AP3B. Use of 700 nm 2P excitation or longer UV (∼360 nm) avoids shorter wavelength DNA absorptions. **C**. Two-photon absorption cross-sections of the AP3B photobiotinylation probe. **D**. The minimal volume (assuming a single XYZ spot) calculated for three different numerical aperture (NA) objectives. Using high NA, it is possible to biotinylate extremely small volumes within the nucleus. **E**. Streptavidin-Alexa 647 labeling of different sized photo-biotinylated regions (1x1, 2x2 and 4x4 pixel ROI box sizes) in a FISH-fixed U2OS nucleus after photo-biotinylation taken at 5 different cross-linking intensities. The 700 nm femtosecond pulses were delivered through 63x/1.4 Zeiss objective lens. Pixel size was 50 nm (5.3x zoom) and the pixel dwell time as 4.1µs and 30 passes over the ROI was used (123 µs per 0.0025 µm^3^) for all crosslinking powers shown. **F**. Lateral and axial dimensions a one-pixel region cross-linked using 6.28 mW at the above scanning parameters. **G**. Overview of the Femto-seq method (see text). **H**. U2OS transgene structure (adapted from ref. 3) used for Femto-seq validation experiments. **I**. U2OS cell after gene activation (+DOX) and IPTG removal to allow YFP-LacI binding for visualization of transgene site showing colocalization of the YFP-LacI and MCP-mCherry.

Figure 1 shows an overview of the method and the possible photo-biotinylation volumes (Fig. 1A-F). We validated Femto-seq with a transgenic U2OS 2-6-3 cell line (transgene shown in Figure 1H) provided by the Spector Laboratory^3^ The cells were further modified by stable co-transfection with pSV2-EYFP-LacI plasmid to express LacI-YFP, and the transgene was located by imaging the LacO/LacI-YFP spots. Cells are maintained in the presence of IPTG until an hour before use to block cell division inhibition by LacI binding to the LacO repeats. The cell line also allows for Tet/Dox inducible transcriptional activation, which can be monitored by CFP expression and/or MCP-RFP localization. Based on initial experiments to determine expected losses during chromatin pull-down and isolation, and the small size of the nuclear volume around the transgene locus being irradiated, we estimated ∼15,000 cells per experiment would be required for obtaining sufficient photobiotinylated DNA for sequencing. Cells were cultured on a coverglass-bottom dish and one hour prior to imaging incubated with the photobiotinylation reagent. The imaging and photoactivation were carried out using a Zeiss LSM 880 multiphoton/confocal microscope. YFP labeled loci were identified by confocal imaging. We created a macro on the LSM 880 to transfer image data to a lab-written dynamic link library (DLL) that found the transgene loci and returned the corresponding ROIs to the Zeiss software. We used the photobleaching module to irradiate the ROIs with 700 nm femtosecond pulses delivered through a 0.5 NA air objective lens. We irradiated a 4x4 pixel box (3.32 x 3.32 µm) centered on the activated transgene. The extent of the axial 2P excitation was ∼2.5 microns, and we estimate the photo-biotinylated volume targeted around the transgene locus to be ∼28 μm^3^. After irradiation of the ROIs, cells were incubated for one hour in 1,6-hexanediol to remove non-covalently linked AP3B from nuclei. To account for bias associated with AP3B chromatin accessibility, ∼15,000 cells were incubated with AP3B as described above and uniformly irradiated by UV light (365 nm) in a StrataLinker 2400 (Stratagene) UV box. Biotinylated DNA from the UV irradiated cells was isolated, sheared, captured with Streptavidin beads, and sequenced and used to normalize coverage. Measurements of U2OS nuclear volume in our cells yielded a value of ∼600 µm^3^ (similar to a value of ∼700 µm^3^ reported in ref. 4). Assuming spatially homogenous chromatin, we would expect an enrichment factor of ∼21 (600/28). After a 1,6-hexanediol wash step to remove non-covalently bound AP3B, purification of the biotinylated DNA fraction and sequencing, a 14.8-fold enrichment of the targeted transgene sequences was found over untargeted rDNA background sequences (Fig. 2A and B) after normalizing crosslinking to the uniformly UV irradiated sample. Given the assumptions made above, this is in reasonable agreement with the predicted enrichment.

**Figure 2.**
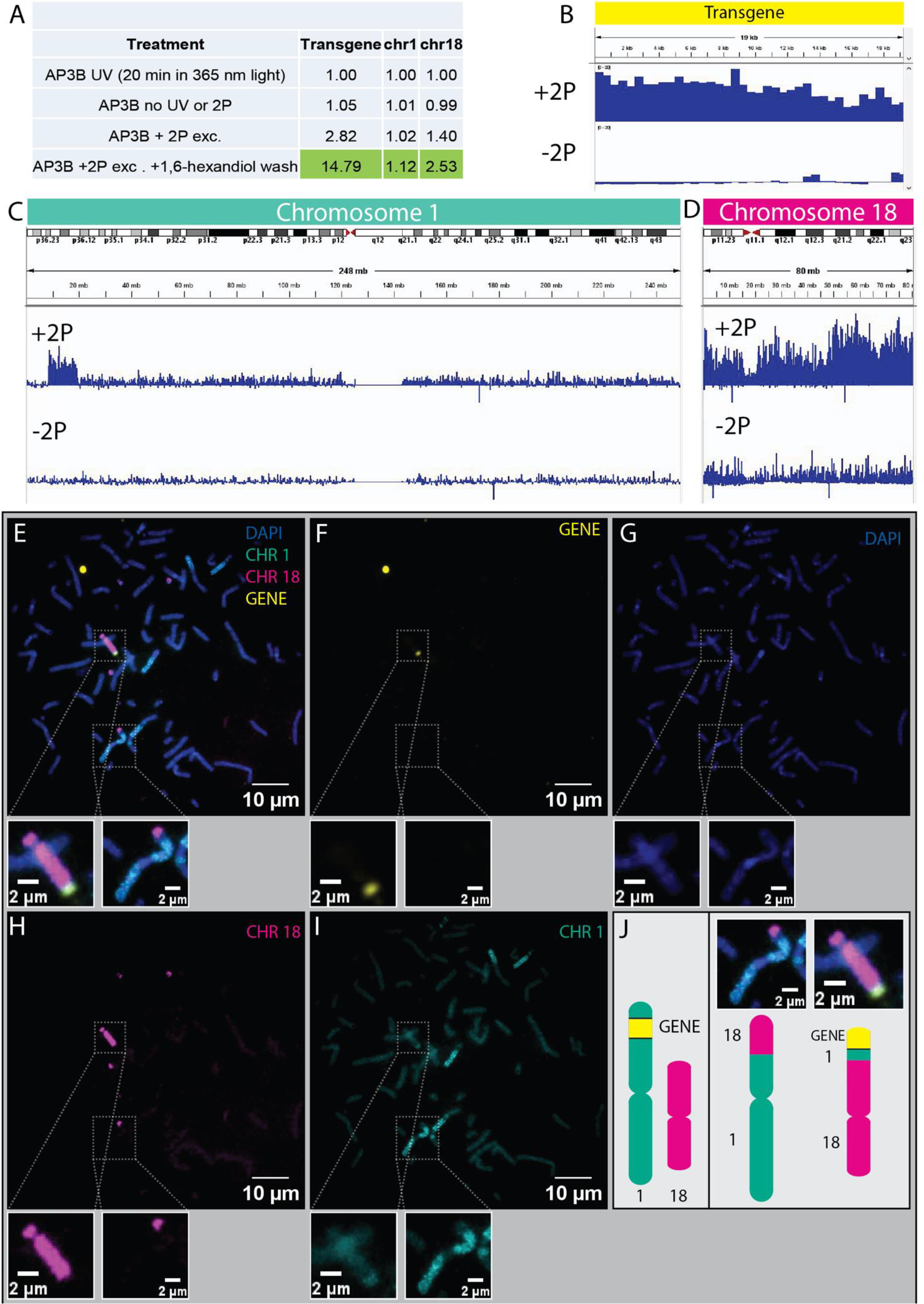
Targeted photo-biotinylation and isolation of chromatin from a transgene locus in cultured U2OS cells. **A**. Calculated enrichments of selected genomic regions from different treatments (normalized to uniform UV irradiation reads). **B-D**. Browser shots of transgene, Chromosome 1, and Chromosome 18, respectively, mapping read coverage as the ratio of targeted 2P-treated to UV-treated to control for non-uniform AP3B selectivity. (Unmapped reads in C are q12 peri-centromeric region.) **E-I**. Metaphase DNA-FISH on U2OS cells. Transgene locus in yellow, Chr1 in green, Chr18 in magenta, DAPI in blue. **J**. Schematic of the presumed transgene location from ref 3 (left) and the translocated transgene (right) which we determined initially with Femto-Seq, and then subsequently verified with multicolor FISH.

Interestingly, a genome-wide analysis found that both chromosomes 1 and 18 had significant enrichment. Coverage plots showed the enrichment of Chr1 was around a 10 MB region of the p36 arm, consistent with the reported insertion site^3^ (Fig. 2C). To explore the appearance of the Chr18 enrichment in our Femto-seq results we performed FISH experiments targeting Chr1 and Chr18 to visualize proximity to the transgene. Metaphase chromosomes (Fig. 2E-I) confirmed that a translocation between the tip of Chr1 where the transgene was originally inserted and Chr18 had occurred, explaining the chromosome-wide enrichment of Chr18 (Fig. 2D), in addition to the expected Chr1 p36 arm sub-chromosomal enrichment.

In our validation experiments we used lightly-permeabilized live cells, but Femto-seq can also be applied to cells prepared for DNA-FISH enabling any gene of interest to be targeted. AP3B will intercalate and can be photo-crosslinked in cells fixed and prepared for FISH, as shown in Figure 1E and in the Supplementary Information section (Figure S1). DNA can also be harvested from these cells as prepared under most FISH protocols (see SI, Figure S2).

Using Femto-seq we can isolate DNA for sequencing from any volume of interest in the nucleus that can be located by high resolution of imaging, for example, to uncover sequence-specific relationships that may exist around nuclear bodies or near active super-enhancers. Because individual cells are imaged, Femto-seq also allows for specific phenotypes within a population to be exclusively targeted. In summary, our method is a unique new tool for analyzing 3D genomic architecture that sits in a unique space between imaging-based methods and chromosome capture techniques.

## Acknowledgements

We thank Dr. Hening Lin and his laboratory for the generous gift of the AP3B reagent. We acknowledge the support for this work from the NIH common fund 4DN Nucleome Project U01-HL129958 to John Lis, Hening Lin, Warren R. Zipfel, and R01-GM145934 to Warren R. Zipfel

## Online methods section

### Photo-biotinylation reagent

Femto-seq utilizes a specifically designed photoactivatable DNA crosslinker with an affinity tag, 4′-Aminomethyltrioxsalen-PEG3-Biotin (AP3B). The photoactivatable component is a psoralen derivative, 4’-aminomethyltrioxsalen (4’-AMT). The compound was synthesized as described in ref 1. The compound has a polyethylene glycol trimer linker to increase solubility. The 3-PEG is terminated with a biotin moiety, which is used as the affinity tag. AP3B is similar to the commercially available EZ-link psoralen-PEG3-biotin probe (ThermoFisher # 29986), but we have found AP3B to be a more efficient photo-crosslinking reagent.

### Cell line modifications

A modified clone of the human bone osteosarcoma epithelial U2OS 2-6-3 cell-line^2^ was gifted by the Spector Laboratory at Cold Spring Harbor Laboratory. The cells feature ∼200 copies of a transgenic construct, each containing 256 Lac Operator (LacO) repeats, 96 tetracycline-responsive elements which regulate a minimal CMV promoter, and an open reading frame. Transcription of the coding sequences produce CFP-tagged SKL proteins, as well as mRNA containing 24 MS2 sequences when activated. We modified the cell line to stably expresses LacI-YFP to locate the transgene locus by confocal microscopy by accumulation of LacI-YFP at the LacO repeats, and rtTA to enable transcription by the addition of doxycycline. This was accomplished by co-transfection with pSV2-EYFP-LacI plasmid and pCDH puromycin resistant plasmid at a ratio of 1:20. 5mM Isopropyl β-D-1-thiogalactopyranoside (IPTG, Sigma-Aldrich I5502) was added to the media to allow cell division, since binding of LacI-YFP to the LacO site will block DNA replication. IPTG was replenished every 2-3 days. 1 μg/ml puromycin was added to the select successfully transfected cells. To add the rtTA construct, U2OS 2-6-3 LacI-EYFP cells were co-transfected with pTetOn plasmid and pCDH blasticidin resistant plasmid at a ratio of 1:20. 15 μg/ml blasticidin was added to the select successfully transfected cells. When the distinct single colonies were formed (2-3 weeks after selection started), 72 were selected and plated onto two 48-well plates (with puromycin and IPTG). 50 cells survived and were plated on MatTek dishes for imaging. We found 9 that showed CFP-SKL expression after adding doxycycline, and 7 monoclonal cell-lines showed bright yellow spots after IPTG removal. The monoclonal cell-lines that showed bright yellow spots in the nuclei and CFP-SKL expression in the cytosol were frozen for later use.

### Cell culture

U2OS 2-6-3 cells were kept in a 37°C incubator in Dulbecco’s modified Eagle’s medium (DMEM) containing 10% fetal bovine serum (FBS) and 1% antibiotic-antimycotic (Gibco). The media, also containing 5μM of IPTG, was replenished every 2 days. Cells were cultured on tissue culture-treated plastic at 37°C and 5% CO_2_, and maintained at 10-80% monolayer confluency.

### Cell sample preparation for Femto-seq

10,000 U2OS 2-6-3 cells stably expressing LacI-YFP and pTetOn were plated at the center of a #1.5 coverslip-bottomed (7 mm dia.) MatTek dish in a 50 μL volume. After allowing the cells to settle on the surface of the plate, 2 mL of DMEM containing 5 mM IPTG and 5 μg/mL doxycycline was added to the dish. Cells were then incubated overnight at 37°C. The following day, 6 hours before irradiation, media was removed and 2 mL of fresh DMEM media without IPTG was added. One hour before irradiation cells were permeabilized by incubation in 0.2% Triton-X-100 in a stabilizing buffer (4% PEG 8000, 1mM EGTA, 10mM PIPES) for 5 minutes. Cells were then washed three times with 1 mL PBS and then incubated in 200 μM AP3B in PBS for 1 hour at 37°C.

### Confocal Imaging, ROI targeting and microscope automation

For the transgene validation experiments reported here, LacI-YFP loci were visualized using a 20x/0.5NA objective lens and 514 nm excitation. Once a field of view was imaged, a Zeiss Zen VBA macro is run which sends the image data to a dynamic link library (DLL) for analysis. The DLL finds the fluorescent loci and exports a text file of the rectangular ROIs around each locus back into the Zen macro. The loci (fluorescent spots) are found using the algorithm described in ref. 3. The user then uses the photobleaching module to irradiate the ROIs with 700 nm two-photon excitation to photobiotinylate the DNA. The macro and DLL are available from the corresponding author.

**Figure S1.**
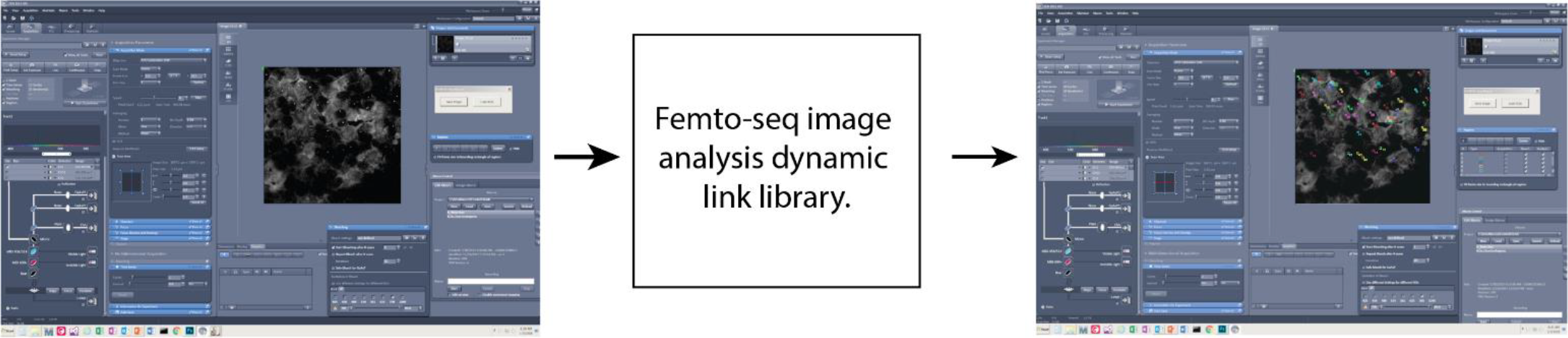
Use of Zeiss LSM 880 confocal/multiphoton system for Femto-seq. A VBA (Visual Basic for Applications) macro written for the Zeiss Zen software (version 2.3 SP1) sends the image data (2D array of bytes - typically 512 x 512) to a dynamic link library (Femtoseq.dll) written in C/C++ (Visual Studio 2019) which locates the fluorescent loci and generates ROIs. The ROIs are imported back into Zen and laser irradiation of the regions using 700 nm femtosecond laser pulses is carried out using the ZEN photobleaching module.

In the validation experiment presented here, 4x4 pixel boxes centered on the LacI-YFP locus were used to specify the regions to irradiate. After a field of view was irradiated, the stage was moved to a new field of cells and the process repeated. Around 5000 cells could be irradiated per dish before the YFP contrast was lost (due to LacI-YFP disassociation). In these experiments 30 passes over each ROI was used. Laser power was ∼70 mW delivered through a 0.5 NA objective lens, with a pixel dwell time of 6.3μs and pixel size of 50 x 50 nm. The power we used to activate the AP3B cross-linker and biotinylate the DNA in the transgene experiments was higher than, for example, the powers used in the power series shown in Figure 1E. Adjusting for the differing numerical apertures (1.4 in Figure 1E vs 0.5 in our transgene experiments) and based on the measured two-photon absorption cross-section at 700 nm (∼1.5 GM), the number of excitations per pixel dwell time at the center of the point spread function would be ∼50 at 0.5 NA and 70 mW, compared to ∼70 using a 1.4 NA objective at 12.5 mW (i.e. as in Figure 1E). For our transgene validation experiments, we used higher powers both due to the lower NA and the fact that we do not know the two-photon *cross-linking* quantum yield. We also were trying to capture a larger volume around the transgene site, so the higher power produced a greater crosslinking depth (i.e. as in Fig. 1E).

### Initial optimization experiments

As we developed the Femto-seq methodology, we carried out a number of initial experiments using the U2OS transgene cells that led to the successful demonstration experiment shown here. These involved different laser powers and ROIs doses, and various “tweaks” to the isolation procedure which we will not cover in detail, with the exception of noting two areas which we have found to be critical. The first is the need for accurate 3D co-alignment of the femtosecond laser focus with the confocal excitation used to locate the target regions. The second is the use of 1,6-hexanediol to help remove non-covalently bound AP3B from the chromatin before streptavidin pulldown. The latter reduced the non-targeted background DNA sequences by more than a factor of 5 (Fig. 2A).

### Femto-seq Library Preparation for Illumina Sequencing

Following irradiation, cells were washed with 1,6-hexanediol (50mM) for 1 hour on the dish. The cells were then detached from the dish with 0.05% Trypsin-EDTA (Gibco) and spun down and stored at -80°C until DNA extraction. Genomic DNA from cross-linked (2P or UV) and non-cross-linked cells were extracted and fragmented to 200-800 bp by sonication. Biotinylated fragments are then captured by Dynabeads MyOne C1 streptavidin beads, and while on beads, end repair, A-tailing, and TRUseq adapter ligation was performed. Adapter ligated fragments are then eluted, and psoralen cross-links chemically reversed in the presence of 3M Urea, 0.1 M KOH, 1 mM EDTA at 90°C. The resulting library is PCR amplified and submitted for paired-end Illumina sequencing (2x 37bp) on a HiSeq 2000 or a NextSeq 500 instrument.

The resulting sequencing data is processed using a custom script. Briefly, paired-end reads are mapped to a custom human genome where the transgene plasmid sequences are included as a separate chromosome using bowtie2 software^4^. Mapped reads were visualized using IGV Genome Browser and reads that mapped to regions of interest were counted using bedtools^5^ and deeptools software^6^.

### Femto-seq Library Preparation Protocol

#### A. DNA isolation

1. 2P- or UV-crosslinked cells (15-30K cells in 300 μl 1xPBS) were lysed with equal volume of 2x Lysis Buffer (200 mM Tris-Cl pH 8.5, 200 mM EDTA, 2% SDS).
2. Incubate at 70°C for 30 min.
3. Cool down to room temperature (∼15 min).
4. Add 5 μl of RNase A/T1 cocktail per sample.
5. Incubate at 37°C for 30 min.
6. Add 86 μl of 8M KAc (final concentration 1M). Mix gently.
7. Incubate on ice for 30 min.
8. Centrifuge at room temperature max speed for 15 min to pellet proteins and cellular debris.
9. Transfer supernatant to a new tube.
10. Extract DNA with equal volume Phenol/Chloroform (1:1) mix. Mix thoroughly but gently.
11. Centrifuge at room temperature max speed for 5 min.
12. Transfer upper aqueous phase to a new tube.
13. Repeat Phenol/Chloroform extraction once more.
14. Extract DNA with equal volume Chloroform. Mix thoroughly but gently.
15. Centrifuge at room temperature max speed for 3 min.
16. Transfer upper aqueous phase to a new tube.
17. To precipitate genomic DNA (∼600 μl), add 2 μl of GlycoBlue co-precipitant, 1/10 volume 3M NaAc pH 5.2, and equal volume of isopropanol.
18. Incubate at room temperature for 30 min.
19. Centrifuge at 4°C max speed for 30 min. Discard supernatant after centrifuge.
20. Rinse the pellet with 750 μl 70% EtOH.
21. Centrifuge at 4°C max speed for 5 min. Discard supernatant after centrifuge.
22. Air dry pellet for ∼5 min.
23. Resuspend each pellet in 200 μl of 1x TE (10 mM Tris-Cl pH 7.5 + 0.1 mM EDTA)
24. Add 1 μl of RNase A/T1 cocktail and incubate at 37°C for 30 min.
25. Fragment the genomic DNA to 200-800 bp by sonication using Diagenode Bioruptor (Denville, NJ) at low setting, 30 sec ON/60 sec OFF for 15 min with ice-cold water in the reservoir.
26. Store DNA at -20°C.

#### B. Biotinylated DNA Pull-down with Streptavidin beads

1. Thaw fragmented genomic DNA on ice.
2. Resuspend Dynabeads MyOne Streptavidin C1 beads by vortexing.
3. Transfer 40 μl of bead slurry per sample to a clean tube.
4. Wash the beads 3 times with 5 volumes of TWB buffer (0.05% Tween-20, 5mM Tris-Cl pH 8.0, 0.5 mM EDTA, 1M NaCl). Use magnet to collect beads on the side of the tube and remove the buffer carefully with a pipette.
5. Resuspend beads with 5 volumes of 2X BWB buffer (10mM Tris-Cl pH 8.0, 1mM EDTA, 2M NaCl).
6. Add equal volume (200 μl) of pre-washed beads to each sample.
7. Incubate at room temperature for 30 min on a rotator (or Thermomixer @600 rpm).
8. After binding wash beads, twice with 300 μl of 1x BWB buffer, and twice with 300 μl TWB buffer.
9. For each wash, collect the beads on the side of the tube using a magnet for 2 minutes, remove and discard the buffer by pipetting. Transfer beads with the new wash buffer to a new tube each time.

#### C. DNA End Repair on Beads

1. Resuspend beads in End Repair mix:

2. Incubate at RT Thermomixer 600 rpm for 30 min.
3. Wash beads 2x 0.4 ml 1x BWB buffer and once with 0.4 ml 1x NEB Buffer 2.
4. Resuspend beads in A-tailing master mix:

|  | Per sample |
| --- | --- |
| DNA on beads |  |
| 10X T4 DNA Ligase Buffer (NEB) | 5 µl |
| 2.5 mM dNTP mix | 5 µl |
| T4 DNA Polymerase | 2 µl |
| T4 PNK | 2 µl |
| Klenow Polymerase | 0.5 µl |
| ddH2O | 35.5 µl |

5. Incubate at 37C Thermomixer 600 rpm for 30 min.
6. Wash beads 2x 0.4ml 1x BWB buffer and once with 0.4 ml 1x homemade T4 DNA Ligase buffer (-ATP).
7. Resuspend beads in 50 μl of 1.1X T4 DNA Ligase Buffer (NEB, with ATP).
8. Add 2 μl of 0.3 uM Indexed TRUseq adapter + 3 μl of T4 DNA Ligase (NEB, 1200 units).
9. Incubate ligation reaction at 16°C overnight.
10. Next day, wash beads 2x 0.4ml 1x TWB buffer and once with 0.4 ml BWB buffer. Transfer beads to a clean tube with each wash.
11. Elute DNA from beads with 56 μl Elution Buffer (10 mM EDTA, 95% Formamide). Incubate at 65°C for 10 minutes.
12. Collect beads on magnet and transfer supernatant to a new tube.
13. Collect residual beads on magnet and transfer supernatant to a new tube again. This step is critical - if any residual magnetic beads are left in the elution it will cause DNA degradation during the next crosslink reversal step.

|  | Per sample |
| --- | --- |
| DNA on beads |  |
| 10X NEB Buffer 2 | 5 µl |
| 1.0 mM dATP | 10 µl |
| Klenow exo-polymerase | 2 µl |
| ddH2O | 33 µl |

#### D. Psoralen crosslink reversal and library prep

Crosslink reversal buffer:

|  | Final conc | 1x |
| --- | --- | --- |
| DNA Eluate | | 56 μl |
| 8mM Urea | 3M | 37.5 μl |
| 2M KOH | 0.1M | 5 μl |
| 50mM EDTA | 1mM | 2 μl |

1. Incubate at 90°C for 10 minutes in the above buffer.
2. Neutralize and EtOH precipitate DNA using:

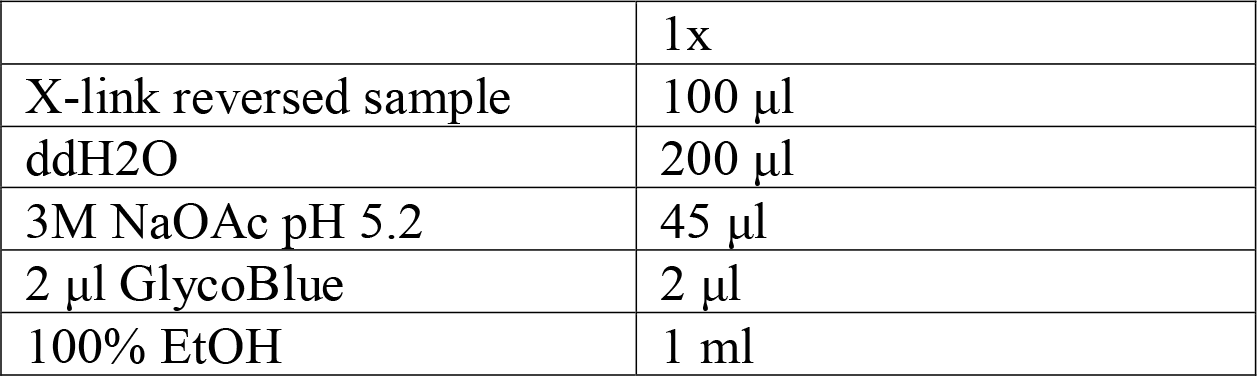
3. Incubate at -20°C for 1 hour.
4. Centrifuge at 4°C max speed for 30 min. Discard supernatant after centrifuge.
5. Rinse the pellet with 750 ul 70% EtOH.
6. Centrifuge at 4°C max speed for 5 min. Discard supernatant after centrifuge.
7. Air dry pellet for ∼5 min at RT.
8. Resuspend each pellet in 50 μl ddH2O.
9. Perform pilot PCR to determine optimal number of amplification cycles:

1. Incubate at 90°C for 10 minutes in the above buffer. 2. Neutralize and EtOH precipitate DNA using:
|  | 1x |
| --- | --- |
| X-link reversed sample | 100 μl |
| ddH<sub>2</sub>O | 200 μl |
| 3M NaOAc pH 5.2 | 45 μl |
| 2 μl GlycoBlue | 2 μl |
| 100% EtOH | 1 ml |

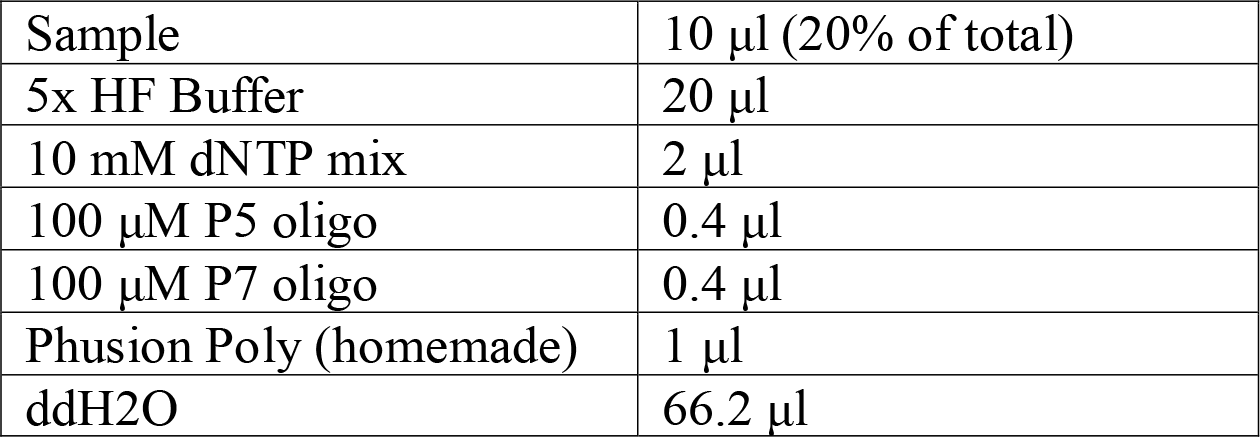
10. Divide each 100 μl PCR reaction into 4x 25 μl aliquots. Run the following PCR with varying cycle numbers (14, 18, 22, 26 cycles). 95°C 5minutes, (95°C 20sec, 58°C 20sec, 72°C 30sec - 14, 18, 22, 26x, 72°C 5min, 4°C ∞
11. Run 12 μl (half) of each PCR reaction on 1.2% Agarose gel and determine the appropriate PCR cycle for the final library amplification.
12. Perform Final PCR amplification for library prep:

13. Divide each 400 ±l PCR reaction into 4x 100 μl aliquots. Run the following PCR with predetermined number of cycles.
14. After PCR, EtOH precipitate PCR products to reduce sample volume for AmPure bead clean-up.
15. Resuspend each sample with 100 μl ddH2O.
16. Add 160 μl AmPure bead slurry (1.6X) to each sample. Incubate at RT for 5 min on rotator (or Thermomixer 600 rpm).
17. Collect beads on a magnet. Discard supernatant.
18. Wash bead pellet 2x with 200 μl 75% EtOH without disturbing the pellet. Discard the wash
19. Air dry beads ∼5min at RT.
20. Elute DNA with 20 μl of 10 mM Tris-Cl pH 8.5 (Qiagen EB buffer).
21. Quantitate DNA concentration using 2 μl samples with Qubit HS dsDNA Assay.
22. Submit samples for Illumina sequencing (ideally paired-end).

|  |  |
| --- | --- |
| Sample | 40 μl (80% of total) |
| 5x HF Buffer | 80 μl |
| 10 mM dNTP mix | 8 μl |
| 100 μM P5 oligo | 1.6 μl |
| 100 μM P7 oligo | 1.6 μl |
| Phusion Poly (homemade) | 4 μl |
| ddH<sub>2</sub>O | 264.8 μl |

### Femto-seq on cells fixed using DNA-FISH protocols

#### Metaphase DNA-FISH

DNA-FISH was performed using commercially available FISH PAINT probes from KromaTiD (Longmont, CO) and performed using their standardized protocol. In short, cells were arrested in metaphase with 10 μl of 10 μg/ml colchicine and cells harvested after 4 hrs. After swelling using 75 mM KCl hypotonic solution, suspensions were fixed in 3:1 methanol and acetic acid. 20-30 μl of cell suspension was dropped onto glass slides and allowed to dry as per standard metaphase slide preparation^7^. Widefield images were taken using a 63x oil 1.4 NA objective on an inverted Nikon eclipse TI fluorescence microscope.

#### DNA-FISH Streptavidin Staining

Alexa-647-Streptavidin staining was performed on Femto-seq photobiotinylated samples to demonstrate the method also works in fixed samples. Samples were both fixed using a 4%PF 3D FISH protocol and 3:1 methanol acetic acid fixation for metaphase preparations. Samples prepared for metaphase FISH were prepared following the manufacturer’s protocol (KromaTiD) as described above. Samples prepared for 3D FISH were prepared following standard protocols^8^. Both types of fixed samples were washed in PBS, then incubated in 200 μM AP3B in PBS for 1 hour at 37°C and immediately used for site specific photobiotinylation. Following photobiotinylation, cells were washed with PBS for 1 hr at 37°C. Then 1 ml of 1:500 dilution of 1 mg/ml Alexa-647-streptavidin was incubated with the cells for 30 minutes. Cells were then washed with PBS for 30 minutes and stained with DAPI mounting solution and imaged.

**Figure S2.**
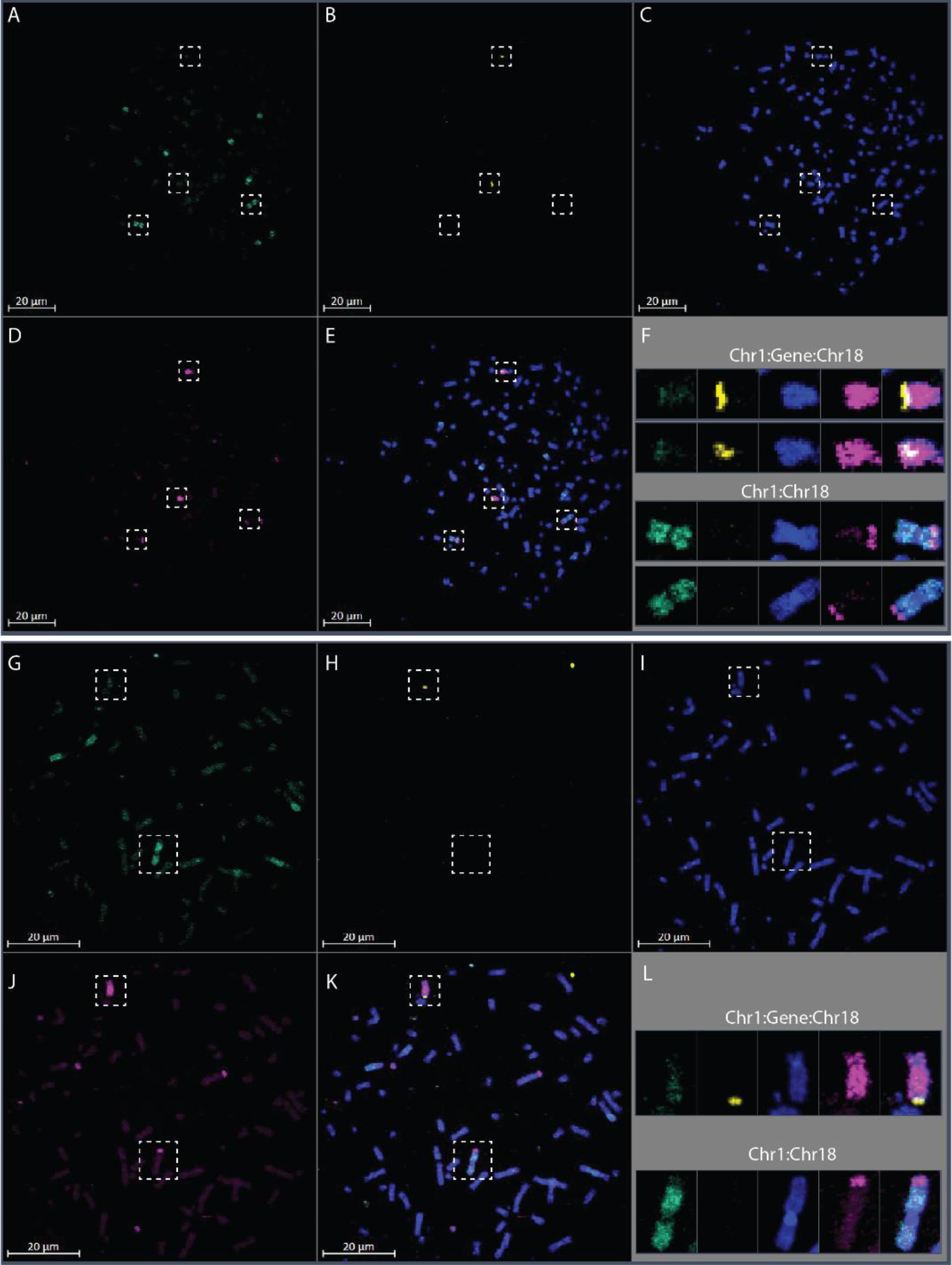
Representative Metaphase Spreads. **A-E** Metaphase DNA-FISH on U2OS cells. LacO locus in yellow, Chr1 in cyan, Chr18 in magenta, DAPI in blue. Boxes around the selected chromosomes show the chr1: chr18 translocation and new location of the YFP locus, **F** Enlarged view of the selected chromosomes showing the translocation of the YFP locus to Chr18. **G-K** Metaphase DNA-FISH on U2OS cells. LacO locus in yellow, Chr1 in cyan, Chr18 in magenta, DAPI in blue. Boxes around the selected chromosomes show the chr1: chr18 translocation and new location of the YFP locus. **L** Enlarged view of the selected chromosomes showing the translocation of the YFP locus to Chr18

**Figure S3.**
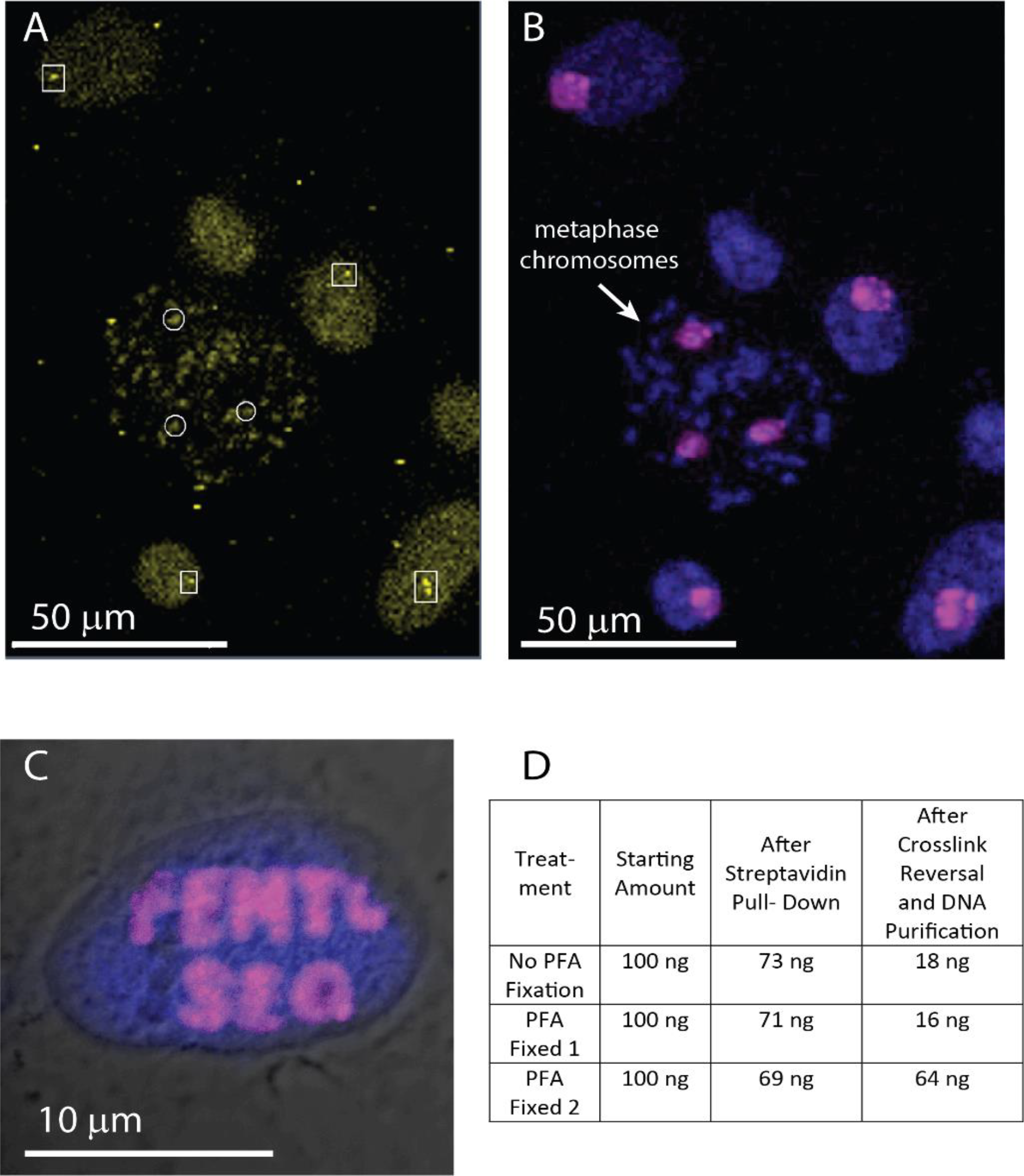
Demonstration of AP3B photobiotinylation in fixed cells. **A**. DNA-FISH to the YFP locus in cells fixed in Carnoy’s fixative (3:1 methanol and acetic acid) for metaphase spread, white boxes show the areas for localized biotinylation using the 700 nm two-photon targeting. **B**. Alexa-647-Streptavidin (Magenta) and DAPI (Blue) labeling after photobiotinylation showing that two-photon mediated AP3B biotinylation can be used to in both cells and metaphase chromosomes in Carnoy’s fixed cells. **C**. Representative photobiotinylation in 4% Paraformaldehyde fixed cells prepared for 3D-FISH protocol using a standard 3D-FISH protocol^4^. **D**. DNA recovery amounts for both non-fixed cells and fixed in 4% PFA cells under untargeted UV box photo-biotinylation. PFA-Fixed 1 and Non-Fixed used the standard ethanol precipitation for DNA purification after crosslink reversal, while PFA-Fixed 2 used a DNA purification column (Zymo Research DNA Clean and Concentrator).

